# *Ixodes scapularis* does not harbor a stable midgut microbiome

**DOI:** 10.1101/198267

**Authors:** Benjamin D. Ross, Beth Hayes, Matthew C. Radey, Xia Lee, Tanya Josek, Jenna Bjork, David Neitzel, Susan Paskewitz, Seemay Chou, Joseph D. Mougous

## Abstract

Hard ticks of the order Ixodidae serve as vectors for numerous human pathogens, including the causative agent of Lyme Disease *Borrelia burgdorferi*. Tick-associated microbes can influence pathogen colonization, offering the potential to inhibit disease transmission through engineering of the tick microbiota. Here, we investigate whether *B. burgdorferi* encounters abundant bacteria within the midgut of wild adult *Ixodes scapularis*, its primary vector. Through the use of controlled sequencing methods and confocal microscopy, we find that the majority of field-collected adult *I. scapularis* harbor limited internal microbial communities that are dominated by endosymbionts. A minority of *I. scapularis* ticks harbor abundant midgut bacteria and lack *B. burgdorferi*. We find that the lack of a stable resident midgut microbiota is not restricted to *I. scapularis* since extension of our studies to *I. pacificus, Amblyomma maculatum*, and *Dermacentor* spp showed similar patterns. Finally, bioinformatic examination of the *B. burgdorferi* genome revealed the absence of genes encoding known interbacterial interaction pathways, a feature unique to the *Borrelia* genus within the phylum *Spirochaetes*. Our results suggest that reduced selective pressure from limited microbial populations within ticks may have facilitated the evolutionary loss of genes encoding interbacterial competition pathways from *Borrelia*.

## Introduction

Vector-borne pathogens infect over one billion people annually and have expanded at an alarming rate in recent years (Harvell et al., 2002; Jones et al., 2013; Keesing et al., 2010; Vora, 2008). Lyme disease, which is caused by the tick-borne bacterial pathogen *Borrelia burgdorferi*, is the fifth-most reported infectious disease in the United States, corresponding to over 90% of vector-borne infections in North America (Kugeler et al., 2015; Radolf et al., 2012). *B. burgdorferi* transits an enzootic cycle between small mammal reservoir hosts and other mammals, vectored by ticks of the genus *Ixodes* (Radolf et al., 2012). Transmission of *B. burgdorferi* to humans via tick bite results in a constellation of inflammatory symptoms requiring an antibiotic treatment regimen for resolution within 2-3 weeks in most cases (Berende et al., 2016; Wormser et al., 2006). While a vaccine targeting *B. burgdorferi* had been approved by the FDA, it was subsequently withdrawn and currently there is an urgent need for new strategies to control tick-borne disease transmission (Nigrovic and Thompson, 2007).

Colonization resistance against pathogens mediated by commensal microorganisms is one such proposed strategy (Buffie and Pamer, 2013; Finney et al., 2015). Studies in mosquitos and tsetse flies, among others, have motivated the characterization of the endogenous microbiota associated with vectors in hopes of identifying means by which pathogen transmission can be interrupted through direct or indirect interactions (Cirimotich et al., 2011a; Cirimotich et al., 2011b; Wang et al., 2017). One mechanism that may govern interactions between pathogens and the microbiota involves direct competition. Many bacteria possess elaborate mechanisms that can mediate interbacterial competition in polymicrobial environments, including specialized pathways such as the type VI secretion system, which delivers toxic effector proteins to target cells (Hibbing et al., 2010; Russell et al., 2014). It is thought that the presence and repertoire of interbacterial systems in a given bacterial genome reflects to a certain degree the selective pressures that organism faces in its natural niche (Zhang et al., 2012). Indeed, these antagonistic systems contribute to bacterial fitness in complex bacterial communities such as the mammalian gut (Kommineni et al., 2015; Verster et al., 2017; Wexler et al., 2016).

The genus *Borrelia* comprises a group of pathogenic bacteria that rely upon hematophagous arthropods for infectious transmission to humans, including ticks of the order *Ixodida* (Radolf et al., 2012). In hard ticks (*Ixodidae)*, surveys of the internal microbiota have to-date identified bacteria that could potentially restrict pathogen transmission (Andreotti et al., 2011; Budachetri et al., 2014; Budachetri et al., 2016; Clay et al., 2008; Clayton et al., 2015; Hawlena et al., 2012; Khoo et al., 2016; Nakao et al., 2013; Narasimhan et al., 2014; Rynkiewicz et al., 2015; Swei and Kwan, 2016; Trout Fryxell and DeBruyn, 2016; van Treuren et al., 2015; Williams-Newkirk et al., 2014; Zolnik et al., 2016). However, significant variation in the diversity and identity of tick-associated bacteria was observed in these studies – rendering the potential for bacterial interference within ticks uncertain. In this study, we aimed to determine whether *I. scapularis* possesses a diverse midgut microbiota that might be encountered by *B. burgdorferi*. Most previous studies of the microbiome of hard ticks have utilized high-throughput sequencing technologies. We sought to augment sequencing with direct measurements of bacterial load and visualization of bacteria within adult and nymphal *I. scapularis* ticks by confocal microscopy. With these complementary approaches, we provide evidence that hard ticks lack a stable midgut microbiota.

## Results

### Borrelia lacks interbacterial effector–immunity genes

Members of the phylum *Spirochaetes* inhabit diverse environments, from the dense and competitive oral cavity and gastrointestinal tract of mammals to the midguts of arthropods like termites and ticks. How members of the phylum *Spirochaetes* engage in interactions with other bacteria is not understood. We therefore sought to characterize the distribution of interbacterial effector and immunity genes in the genomes of *Spirochaetes*. We first compiled a database of *Spirochaetes* genomes, encompassing a total of 63 genomes representing all major genera. These genomes were queried for the presence of homologs of 220 interbacterial effector and immunity genes (Zhang et al., 2012), including those with characterized domains found associated with contact-dependent inhibition (Hayes et al., 2010), type VI secretion system (Russell et al., 2014), and ESX/T7SS antagonistic pathways (Cao et al., 2016; Whitney et al., 2017). This analysis revealed the presence of genes encoding interbacterial effector and immunity domains throughout the *Spirochaetes*, particularly in species known to inhabit polymicrobial environments such as the mammalian gut microbiome (*Treponema succinifaciens* and *Brachyspira* spp) and the oral microbiome *(Treponema denticola)* (Figure 1A). We failed to detect effector gene homologs and identified only a limited group of immunity gene homologs encoded by any species within the genus *Borrelia*, including representatives from the *sensu stricto* and *sensu lato* genospecies (Becker et al., 2016). Since the phylum *Spirochaetes* is considered to be monophyletic, with extant genera descending from a single common ancestor (Paster et al., 1991), parsimony supports the conclusion that *Borrelia* lost interbacterial effector genes early in the evolution of the genus. Parallel investigation of the genomes of tick-associated endosymbionts and tick-transmitted intracellular pathogens including *Rickettsia*, *Coxiella*, *Anaplasma*, and *Ehrlichia* revealed that these genomes also largely lack interbacterial effector– immunity genes (Figure 1B).

**Figure 1.**
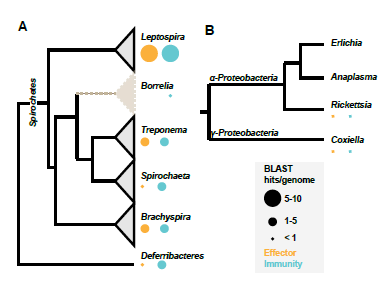
Distribution of interbacterial effector–immunity gene homologues across the phylum *Spirochaetes*. **(A)** The distribution of interbacterial effector and immunity gene homologs in genomes from across a phylogenetic tree of genera within the phylum *Spirochetes*. *Deferribacteres* is shown as the outgroup. BLAST hits are normalized to the number of genomes analyzed for each genus. *Borrelia* shows a significant deviation from the expected frequency relative to other genera (*p* < 0.05). **(B)** Interbacterial effector and immunity gene homologs in genomes of endosymbionts and pathogens associated with *Ixodidae* from the phylum *Proteobacteria*.

### Assessment of bacterial abundance and diversity in I. scapularis

The evolutionary loss of effector–immunity genes from the genus *Borrelia* led us to hypothesize that interbacterial interactions might be limited within *I. scapularis*. We therefore sought to quantify the microbial communities associated with wild *I. scapularis*. We first isolated DNA from the dissected viscera (a combination of internal tissues that included midgut, reproductive tissues, and salivary glands) of 61 adult ticks collected from 5 distinct geographic sites in the Midwestern US. We then performed quantitative polymerase chain reaction (qPCR) targeting conserved regions of the 16S rRNA gene. Our analysis demonstrated that the mean internal bacterial load of unfed adult *I. scapularis* ticks varies over several orders of magnitude (Figure 2A). Male ticks analyzed harbored less bacterial load than did females (4.5x10^5^ vs 5.3x10^7^, t-test p-value < 0.001, N = 61). qPCR with primers specific to the *B. burgdorferi flaB* gene revealed infection frequencies similar to that previously reported from the Midwestern geographic regions (Figure S1A) (Hamer et al., 2014). To investigate the differences in bacterial community underlying the sex-specific differences that we observed, we performed 16S rRNA gene sequencing on the same samples used for qPCR analysis. In addition, we sequenced the external washes from a subset of ticks, and a water-only control for each group. 178 OTUs were detected in the water-only controls, including taxa commonly implicated in reagent contamination and some previously reported to be associated with ticks, such as *Sphingomonas* and *Comamonas* (Nakao et al., 2013; Narasimhan et al., 2014; van Treuren et al., 2015). These OTUs were subsequently eliminated from all other samples in downstream analyses. To describe the taxa 6 that comprise *I. scapularis*-associated bacterial communities following removal of water-borne contamination, we calculated the taxonomic relative abundance averaged across all internal and external samples. This revealed that 258 (95%) of 270 OTUs fell below 1% relative abundance when averaged. In contrast, other taxa exceeded 1% average relative abundance in both internal and external samples, including *Bacillus, Pseudomonas*, and *Enterobacteriaceae* (Figure S2). Importantly, the taxon that was most abundant in internal samples but entirely absent in external samples was the genus *Rickettsia* (average relative abundance of 52%), which includes the dominant *Rickettsia* endosymbiont of *I. scapularis* (Gillespie et al., 2012). The relative abundance of *Rickettsia* exhibited a positive correlation with total bacterial load (Figure 2B), indicating that *Rickettsia* is the primary driver of bacterial abundance in most *I. scapularis* ticks. The family *Spirochaetaceae* also was often abundant in viscera samples but absent in washes (Figure S2).

**Figure 2.**
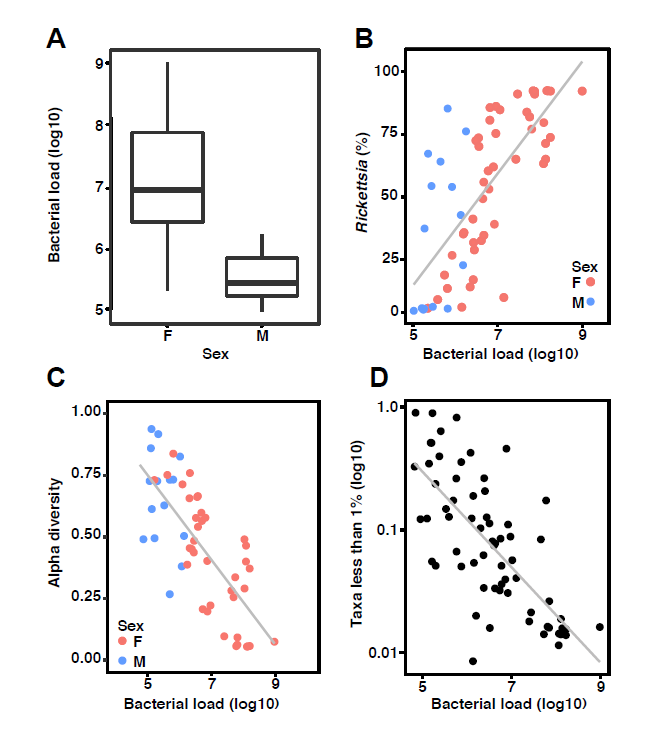
Inflation of bacterial diversity in *Ixodes scapularis*. **(A)** The bacterial load of male and female *I. scapularis* adult ticks, determined by qPCR amplifying the 16S rRNA gene (N=61). Male ticks harbor significantly less bacteria than do females, (t-test *p* < 0.001). **(B)** Correlation between the relative abundance of *Rickettsia* and the total bacterial load across all adult *I. scapularis* ticks in unfiltered samples (Spearman’s rho = 0.75, *p* <0.001). **(C)** Alpha diversity measured by the Simpson’s value (log2) negatively correlates with bacterial load (log10) across all adult *I. scapularis* samples without filtering of low abundance taxa (Spearman’s rho= −0.74, *p* <0.001). Male (blue) and female (pink) samples appear stratified according to both load and diversity. **(D)** Relative abundance of the sum of all taxa whose average relative abundance was less than 1% correlates negatively with bacterial load in adult *I. scapularis* samples (log10), Spearman’s rho = −0.73, *p* < 0.001.

The number of taxa with low relative abundance could indicate that the internal environment of *I. scapularis* is particularly well suited to fostering diverse microbial communities. However, an alternate possibility is that low abundance OTUs represent signal from contamination. Indeed, several studies have examined the effect of low biomass input on diversity metrics in 16S rRNA gene sequencing surveys, with the notable result that alpha diversity (within-sample diversity) correlated negatively with the abundance of input DNA (Jervis-Bardy et al., 2015; Lauder et al., 2016; Salter et al., 2014). We therefore examined the relationship between alpha diversity and bacterial load as quantified by qPCR in our *I. scapularis* samples. When examined across all 61 *I. scapularis* samples, we found a strong negative correlation between total bacterial load and alpha diversity that was independent of geographic origin (Spearman’s rho = −0.74, *p* < 0.001) (Figure 2C and S3). These data imply that the low bacterial biomass associated with adult *I. scapularis* ticks can impact the interpretation of 16S rRNA gene sequencing surveys by artificially inflating alpha diversity. We further observed a strong negative correlation between bacterial load and the relative abundance of the group of taxa contributing to 1% or less of the total across all *I. scapularis* samples (Figure 2D). We found only six taxa to be present in all *I. scapularis* viscera samples, represented by eight OTUs, including the genera *Rickettsia, Borrelia, Pseudomonas, Francisella*, and *Escherichia*, and the family *Enterobacteriaceae* (Figure S2). Linear discriminant analysis effect size (LEfSe) revealed the taxa most likely to explain differences between external wash samples and internal viscera samples. External samples are characterized by taxa from the phyla *Proteobacteria* and *Actinobacteria*, while viscera samples are distinguished by the orders *Spirochaetales* and *Rickettsiales* (Figure S4) (Segata et al., 2011).

While the internal bacteria associated with most wild adult *I. scapularis* are dominated by *Rickettsia* and *B. burgdorferi*, a minority of samples exhibit high relative abundances of three putatively environmental taxa, including the genera *Bacillus* and *Pseudomonas*, and the family *Enterobacteriaceae* (Figure 3A) all of which have been previously described as associated with *I. scapularis* (Moreno et al., 2006; Steinhaus, 1946; van Treuren et al., 2015). Colonization of ticks appeared to be independent for each taxon since, although co-occurrence was detected in a few samples, most ticks harbored only one dominant environmental taxon. While total bacterial load was not correlated with colonization by environmental bacteria, samples with high abundance of these bacteria – deriving from each of the five geographically-isolated collection sites – were less frequently infected by *B. burgdorferi* than expected (Figure 3B, C). This observation suggests that colonization by *Bacillus, Pseudomonas*, and *Enterobacteriaceae* may limit *B. burgdorferi* infection.

**Figure 3.**
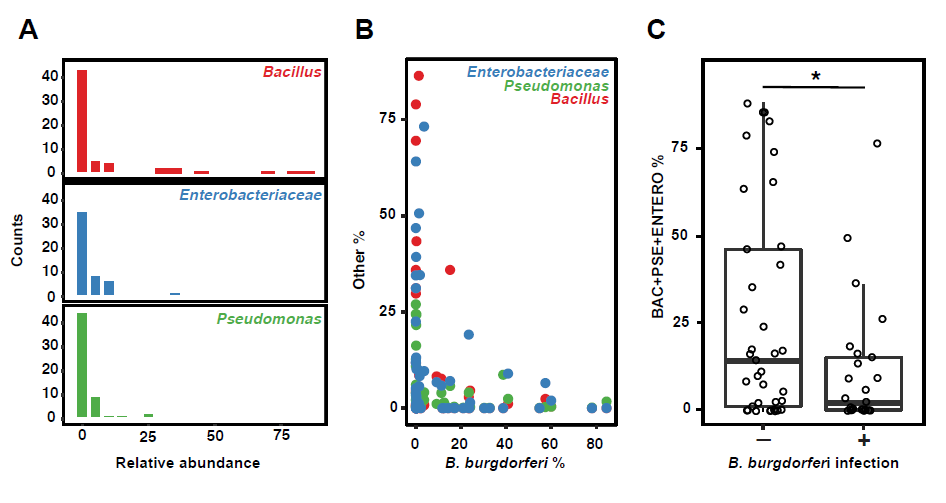
Transient environmental bacteria limit *B. burgdorferi* infection. **A)** Histograms depict the relative abundance of bacteria (*Bacillus*, red; *Enterobacteriaceae*, blue; *Pseudomonas*, green) detected in *I. scapularis* viscera samples, in bins of 5%. Counts indicate number of samples. **B)** Scatter plot of the relationship between the relative abundance of environmental bacteria detected in *I. scapularis* viscera and the relative abundance of *B. burgdorferi*. Color scheme as in (A). **C)** Barplots indicate the sum of the relative abundance for each of three abundant taxa between samples in which *B. burgdorferi* relative abundance exceeds 1% (infected) and those samples for which abundance is less than 1% (uninfected). The difference is statistically significant (Mann Whitney U test, *p* < 0.001).

### Immuno-staining and confocal microscopy reveals I. scapularis midgut biogeography

To orthogonally and directly validate our findings of a limited internal microbiome in ticks, we turned to confocal microscopy in order to visualize bacteria distribution and localization within *I. scapularis*. Our sequencing results suggest that *Rickettsia* comprise the dominant internal microbial inhabitants of most wild adult *I. scapularis* ticks. However, we questioned i) if our analyses failed to detect low-abundance yet highly diverse communities of midgut-associated bacteria masked by the high abundance of *Rickettsia* and ii) if the limited midgut colonization by environmental bacteria that we observed could be visualized and therefore validated. Ticks present a number of challenges that hinder standard microscopy techniques. These challenges include the physical barrier of the thick outer cuticle, a high degree of internal auto-fluorescence derived from the cuticle, and remnants of previous blood-meals including heme crystals (Sojka et al., 2013). In order to preserve tissue integrity and organization, we utilized formalin-fixation and paraffin embedding of dissected ticks, followed by thin sectioning and staining for visualization. Use of biotin-labeled ISH probes allowed us to perform tyramide-signal amplification (TSA), dramatically increasing the signal to noise ratio within tick sections.

With these methods, we characterized the distribution of bacteria within tissues of individual adult *I. scapularis* ticks (N=41). Abundant and DAPI-intense cocci that co-stained with probes targeting universally conserved 16S rRNA sequences (EU338) were observed in the cytoplasm of ovarian cells of all female *I. scapularis* ticks (Figure S5). These are presumed to be vertically transmitted *Rickettsia* (Noda et al., 1997). In striking contrast to ovaries, the midgut of most unfed adult *I. scapularis* lacked bacilli or cocci as visualized by DAPI signal or ISH signal localized to the lumen or luminal epithelium (Figure 5A, 90%, N=41 total ticks). This differed for *B. burgdorferi*-infected *I. scapularis* ticks, in which spirochete cells constituted the only detected midgut bacteria (Figure 4A, S5). However, a minority of *I. scapularis* ticks from a single site (CA) exhibited midgut colonization by bacterial cells with distinct cocci and bacilli cellular morphologies (10% of total *I. scapularis* examined, N=41) (Figure 4B). Since *I. scapularis* nymphs have been reported to possess a diverse midgut biofilm-forming microbiome, with very low relative abundance of *Rickettsia* (Abraham et al., 2017; Narasimhan et al., 2014), we considered the possibility that while adult *I. scapularis* ticks may lack a diverse internal microbiome, earlier life stages could be colonized to a greater extent. To test this possibility, we imaged sections prepared from whole-mounted *I. scapularis* nymphs, using similar formalin fixation and staining methodology as before. We found that *I. scapularis* nymphs also exhibit low DAPI and ISH probe staining within midgut tissues (Figure S6). Since nymphs are considerably smaller than adults, they may be more sensitive to artefactual noise associated with low-biomass high-throughput sequencing. In support of this notion, the alpha diversity of the *Ixodes pacificus* microbiome negatively correlates with life stage progression from larvae to adults as body size increases (Swei and Kwan, 2016).

**Figure 4.**
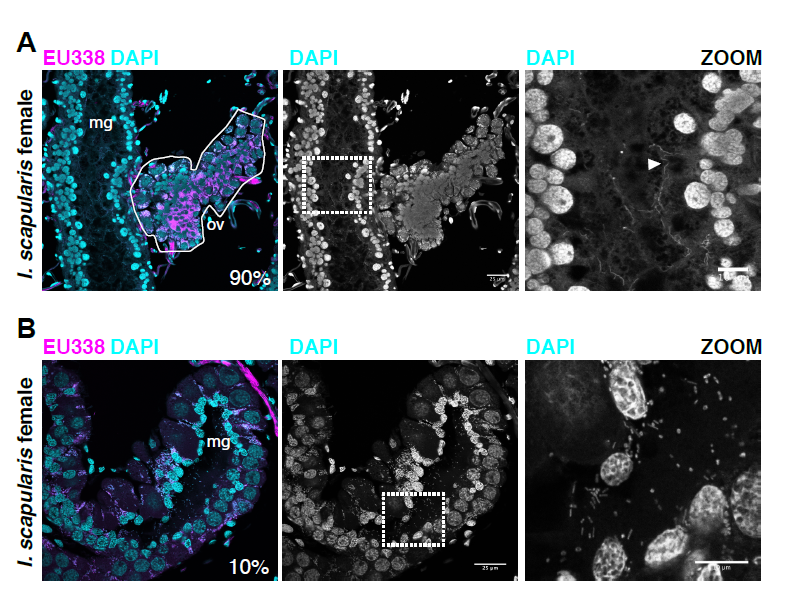
Midgut-associated bacteria in *I. scapularis*. **(A)** Cross-section of the ovary and midgut of a *B. burgdorferi*-infected adult female *I. scapularis* tick. Region highlighted by zoom indicated by white box. “mg” indicates midgut lumen. Ovary indicated by white outline labeled with “ov”. Percent indicates the proportion of analyzed ticks with similar internal bacterial biogeography (N=41). Arrowhead in zoom indicates *B. burgdorferi* cells. **(B)** Staining of an *I. scapularis* female tick collected from CA site with midgut-associated bacteria. White dashed box indicates the region highlighted by zoom. Percent indicates the percent of total ticks analyzed with similar internal bacterial content. Scale bars indicate 25 microns for wide view and 10 microns for zoomed panels.

### 16S rRNA gene sequencing of diverse wild adult ticks

We next sought to test whether our findings of a limited internal bacterial load and restricted diversity in *I. scapularis* were generalizable across *Ixodidae*. Vector competence for *Borrelia* spp. varies across genera within the *Ixodidae*. Ixodid ticks also vector pathogenic *Anaplasma* and *Ehrlichia* spp which must transit the midgut during their enzootic cycles (Ismail et al., 2010). We collected six species of hard tick from multiple geographic locations, including species from the genera *Amblyomma, Dermacentor*, and *Ixodes*. qPCR-based assessment of internal bacterial load revealed a variability similar to that of *I. scapularis*, with *Amblyomma maculatum* exhibiting the highest load and broadest variation across individual ticks (Figure 5A). As observed for *I. scapularis*, the dominant bacterial taxa associated with each tick species were species-specific endosymbionts (Figure 5B). These include *Francisella* and *Rickettsia* in *Amblyomma maculatum, Francisella* in *Dermacentor* spp, and *Rickettsia* in *Ixodes* spp, as has been previously reported (Budachetri et al., 2014; Rynkiewicz et al., 2015). Although alpha diversity varies broadly across species (Figure 5C), negative correlations with total bacterial load across hard tick samples (an exception being *I. pacificus*) supported a similar pattern as seen for *I. scapularis* (Figure 5D).

**Figure 5.**
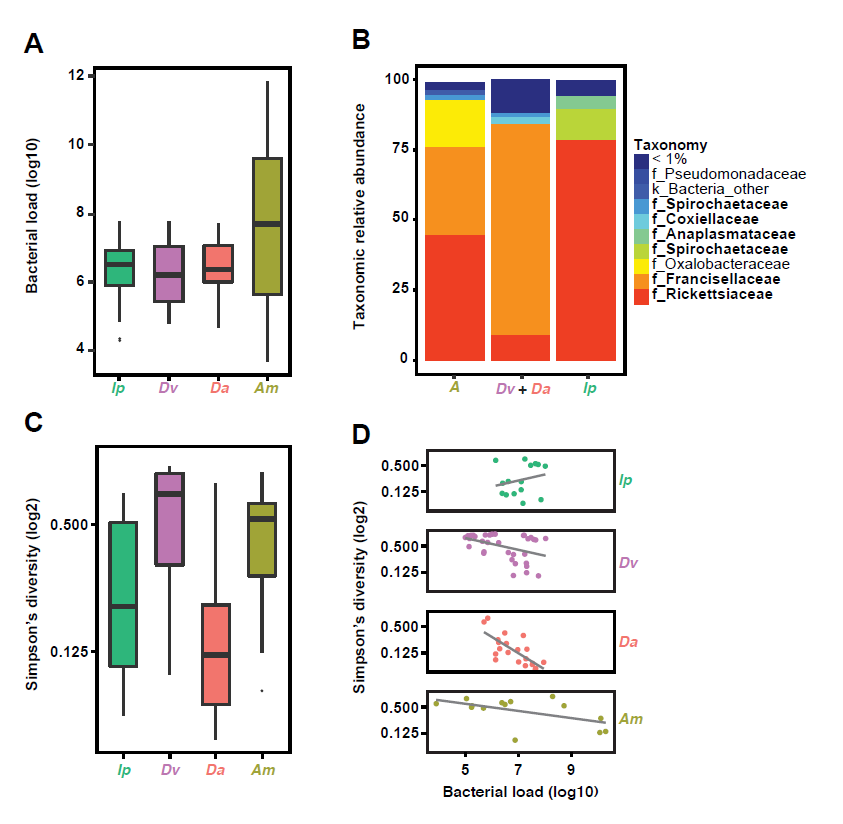
Impact of limited bacterial load on bacterial diversity estimates in hard ticks. **(A)** Box and whisker plots describing the internal bacterial load (log10) across all hard ticks sampled (N=139), as determined by qPCR. Species names abbreviated to *Ip* (*Ixodes pacificus*), *Dv* (*Dermacentor variabilis*), *Da* (*Dermacentor andersonii*), and *Am* (*Amblyomma maculatum*). **(B)** Relative taxonomic abundance for bacteria detected by 16S rRNA gene sequencing averaged across all samples for *A. maculatum*, *D. variabilis* and *D. andersonii*, and *I. pacificus*. Taxa previously known to be internally associated with these species are bolded in legend. **(C)** Box and whisker plots describing Simpson’s Diversity (log2) of all hard tick internal samples without filtering, with same color scheme as in A**. (D)** Bacterial load plotted against Simpson’s Diversity (log2). Spearman’s rank correlation: Ip rho = 0.07, p > 0.1; Dv rho = −0.88, *p* < 0.001; Da rho = −0.73, *p* <0.001; Am rho = −0.62, *p* < 0.05.

Beta diversity (between-samples) analysis using the weighted UniFrac metric (which accounts for differences in taxon abundance between samples) revealed significant clustering of samples by tick genus (ADONIS, p=0.001) (Figure S7A), as expected for stable microbiomes or those largely composed of endosymbionts. In contrast, external wash samples cluster together regardless of which tick they were associated with. This suggests that external microbiomes share common features (eg. the presence of diverse low-abundance taxa and lack of highly abundant endosymbionts) that differ from internal microbiomes of ticks. This interpretation is supported by beta diversity analyses using the unweighted UniFrac metric (in which each OTU is given equal weight regardless of relative abundance in the sample) which revealed no such internal-external (Figure S7B) or genus-level clustering, suggesting that the low-abundance taxa in samples do not sufficiently drive differences between tick genera.

Finally, we sought to extend our microscopy findings across hard tick species. Similar to *I. scapularis*, we found that *A. maculatum, D. variabilis*, and *I. pacificus* ticks lack abundant midgut-associated bacteria, while females belonging to these species contained abundant DAPI-intense cocci within ovaries (Figure S8). We also observed that midgut sections near ovaries sometimes contained localized micro-colonies, similar to those shown to be *Rickettsia* in *I. scapularis*. These may be species-specific endosymbionts, as previously reported (Clay et al., 2008; Klyachko et al., 2007).

## Discussion

Here we have provided multiple lines of evidence showing that unfed wild hard ticks possess a limited internal microbiome. This is in contrast to reports that possession of a diverse microbiome is a characteristic feature of hard ticks across species and life stages (Abraham et al., 2017; Andreotti et al., 2011; Budachetri et al., 2014; Budachetri et al., 2016; Nakao et al., 2013; Narasimhan et al., 2014; Swei and Kwan, 2016; Trout Fryxell and DeBruyn, 2016; van Treuren et al., 2015; Zolnik et al., 2016). Our report adds to the growing literature suggesting that not all animals are intimately associated with a complex and abundant internal microbiome (Hammer et al., 2017; Jing et al., 2014). We find that measurements of high diversity derived from high-throughput sequencing of tick samples is largely a function of low bacterial biomass. This is a widespread problem inherent in high-throughput sequencing studies that rely upon PCR-amplification during the sample preparation process (Glassing et al., 2016; Kim et al., 2017). We propose that pooling multiple samples to increase input biomass is one method that might lower noise from reagent-based or external contamination (Gall et al., 2016).

There is great interest in exploiting natural microbial communities to benefit human health and combat disease. One such avenue of active research involves leveraging the native microbial communities associated with medically-relevant arthropod vectors (like mosquitos and ticks) to prevent pathogen transmission (Cirimotich et al., 2011b; Narasimhan and Fikrig, 2015).

We note that *B. burgdorferi* may be particularly susceptible to inhibition by competitor bacteria if encountered due to a paucity of genetic mechanisms for direct interbacterial interactions. This is supported by our findings that the increased abundance of *Bacillus, Enterobacteriaceae*, and *Pseudomonas* within the midgut is associated with decreased *B. burgdorferi* infection. Notably, these bacteria were also detected in external wash samples. This intriguing pattern suggests that some tick-associated external bacteria can colonize the *I. scapularis* midgut and, in doing so, might competitively exclude *B. burgdorferi*, either by displacement or by inhibition of infection during a subsequent blood meal. Members of these taxa encode an arsenal of mechanisms with which to compete with other bacterial cells, including type IV, VI, and VII secretion systems, contact-dependent inhibition (CDI) mechanisms, and the ability to produce bacteriocins (Cao et al., 2016; Hayes et al., 2010; Riley and Wertz, 2002; Russell et al., 2014; Souza et al., 2015). It is unknown if environmental bacteria can persist within the midgut during transstadial molts, but it is worth noting that detection of *Bacillus* and other bacteria within ticks long preceded the 16S rRNA gene sequencing era (Steinhaus, 1946). Direct inhibition of *B. burgdorferi* by tick-associated bacteria remains an active area of investigation by our laboratories.

Despite the detection of environmental bacteria within ticks, our findings suggest that the midgut of hard ticks may largely be an environment that is ill-suited for bacterial growth, perhaps due in part to the exceedingly low levels of the vital nutrient thiamin in *I. scapularis* (Zhang et al., 2016). While *B. burgdorferi* has evolved unique metabolic strategies in order to persist in this thiamin-limited environment, other less-specialized bacteria may not be able to do so. Additional factors that might limit the overall bacterial load of hard ticks are the possession of conserved or unique innate immunity factors that could target bacteria (Chou et al., 2014; Palmer and Jiggins, 2015), heme toxicity (Anzaldi and Skaar, 2010), and the effects of nutrient limitation and desiccation during the extended period between blood meals (Radolf et al., 2012; Sonenshine and Roe, 2014).

*Borrelia burgdorferi* may encounter a limited diversity of bacteria within the *I. scapularis* midgut, yet co-infections with other pathogens such as *Anaplasma* are common (Hamer et al., 2014). The mechanisms by which *B. burgdorferi* co-exists with other microbial pathogens within the tick remain unexplored. While *Anaplasma, Ehrlichia*, and *Rickettsia* spp are intracellular and may not engage in direct physical interactions with *Borrelia* spp, these bacteria may influence colonization through other means (Kocan et al., 2010; Simhadri et al., 2017). *B. burgdorferi* may not frequently encounter diverse bacteria within the midgut of the tick, however, multi-strain infections may be common within single ticks (Durand et al., 2017; Durand et al., 2015; Herrmann et al., 2013; Strandh and Råberg, 2015; Voordouw, 2015). Furthermore, it is as yet unclear the extent to which different *B. burgdorferi* strains engage in interactions or how multi-strain infections might influence transmission to humans and pathogenicity.

Our bioinformatic analyses suggests that the genus *Borrelia* lost genes for mediating interbacterial interactions during the course of its evolution. While *Ixodes* are the only vectors for *B. burgdorferi* in North America, ticks of the genus *Ornithodoros* (soft ticks) vector the relapsing fever spirochete *B. hermsii* (Dworkin et al., 2002). We did not detect interbacterial effector–immunity gene homologues encoded by *B. hermsii*. Although soft ticks were not included in our 16S rRNA gene sequencing or microscopy, we speculate that *Ornithodoros*, like *Ixodidae*, harbors a limited internal microbiome. Our results further imply that lice of the order *Phthiraptera* (class Insecta), which vectors *B. duttonii* and *B. recurrentis*, may also possess a limited internal microbiome (Lescot et al., 2008). *Borrelia* spp may have therefore evolved to exploit evolutionarily divergent hematophagous arthropods that share the common lack of a stable midgut microbiota.

## Methods

### Tick collection

Ticks were collected from the following sites: oak woodland habitat in Klickitat River Canyon, Washington (*Ixodes pacificus* and *Dermacentor andersonii*); oak-hickory forest in Wolf Creek State Park, Illinois (*Ixodes scapularis* and *Dermacentor variabilis*); oak-dominated forest in Gordie Mikkelson Wildlife Management Area and Carlos Avery Wildlife Management Area, Minnesota (abbreviated GM and CA, *Ixodes scapularis* and *Dermacentor variabilis*); red pine forest in Kettle Moraine State Forest Southern Unit (KM), mixed hardwood forest in Big Eau Pleine County Park (BEP), and oak-hickory forest in Sandberg Woods Conservancy (SC), Wisconsin (*Ixodes scapularis*); oak forest near McPherson Preserve, Oklahoma (*Amblyomma maculatum*). Ticks were collected by the flagging and dragging methods and shipped in 50mL Falcon tubes with damp paper towels on wet ice to retain moisture. Species, developmental stage, and sex of individual ticks were determined by visual inspection.

### DNA isolation

Live ticks were washed three times with sterile water then dried before immobilization on a glass slide using double-sided tape. 21-gauge needles were used to remove the cuticle, and sterile forceps used to dissect and remove the viscera into 500uL sterile deionized water for subsequent DNA purification. Fresh needles were exchanged, and forceps sterilized by washing three times in sterile water and 70% ethanol between each dissection. The first external wash sample was saved, and pooled across individuals of the same species, sex, and geographical collection location. Samples were frozen at −80^°^C prior to DNA isolation. Frozen tissue samples were thawed and subsequently homogenized via glass bead-beating on a MiniBeadBeater (Biospec Products) for two cycles of one minute duration each. Subsequently, phenol-chloroform-isoamyl alcohol extraction of total nucleic acids was performed with a RNase treatment to remove RNA. Following extraction, DNA pellets were resuspended in 25uL sterile deionized water.

### Quantitative PCR

The abundance of bacteria in single tick samples was determined by quantitative PCR using SsoAdvanced Universal SYBR Green Supermix (Biorad) on a BioRad CFX96 Real Time System C1000. Primers used targeted the 16S rRNA gene (331f and 797r(Nadkarni et al., 2002)) or the *B. burgdorferi flaB* gene (Jewett et al., 2007). Standard curves were generated from serial dilutions of purified genomic DNA prepared from monocultures of *Escherichia coli* DH5a or *B. burgdorferi* B31A3. Samples were run in technical triplicates with the mean of each triplicate used for later analysis.

### 16S rRNA gene sequencing and analysis

Genomic DNA from tick samples, external washes, and water controls was submitted for sequencing and individually barcoded for high-throughput sequencing of V3-V4 16S rRNA amplicons on an Illumina MiSeq, in three separate runs (performed by MrDNA). Sequencing reads were subsequently demultiplexed and merged into a single fastq file. The UPARSE pipeline was used to process the samples using default settings (Edgar, 2010, 2013). Following taxonomy prediction and OTU assignment, the OTU table were filtered to only retain OTUs that appeared at greater than 1% relative abundance in at least one sample. Alpha-diversity and beta-diversity metrics were calculated in QIIME (MacQIIME v1.9.0) (Caporaso et al., 2010). For determination of *B. burgdorferi* infection status, tick samples in which the sum of the relative abundances of the family *Spirochaetaceae* exceeded 1% were considered to be infected, while those less than 1% were considered to be uninfected. The sum of the relative abundance of the genera *Bacillus* and *Pseudomonas* was calculated for samples for comparison of *B. burgdorferi* infection status. For LEfSe analysis, the OTU table was converted to relative abundances and, along with associated metadata, was uploaded to the Huttenhower Lab Galaxy web application. Statistical tests and generation of plots were performed in R version 3.3.2 (R Development Core Team, 2013).

### Histology

#### Preparing tick sections

Ticks were washed three times in sterile water and immobilized with double-sided tape attached to glass slides. Cuticles were removed with sterile 21-gauge needles. Following removal of the cuticle, dissected ticks were fixed in formalin for at least 5 days at room temperature. Fixed ticks were then embedded in paraffin and four 4-micron sections were prepared from each tick. Slides were deparaffinized in a series of washes in xylenes, followed by rehydration in washes of decreasing percent ethanol solutions.

#### Antibody-based microscopy

Deparaffinized slides were first subjected to a 40 minute incubation in sodium citrate buffer at 95^o^C in a vegetable steamer (Black and Decker). Slides were then washed in 2X SSC + 0.3% Tween 20, blocked in PBS + 5% BSA, and incubated with BacTrace fluorescein isothiocyanate goat anti-*Borrelia* whole cell antibodies (Kirkegard and Perry Laboratories) for 1 hour at room temperature or overnight at 4^o^C. Slides were then washed, dried, and mounted in ProLong Diamond plus DAPI (Thermo Fisher Scientific). Coverslips were sealed with nail polish.

#### In situ hybridization analysis

Deparaffinized slides were washed in 2X SSC + 0.3% Tween20 then were lightly digested with Proteinase K to allow probe access into tissues, washed and dried. Probes used were EU338 (Amann et al., 1995) or specific to *Rickettsia*. For Rickettsia-specific probes, the sequence of the Rick_1442 16S rRNA probe (Vannini et al., 2005) was aligned to the publically available REIS genome (Gillespie et al., 2012) and confirmed to be identical. ISH-probes (5’ biotin labeled) were diluted 1:100 to 50uM in ethylene carbonate hybridization buffer (15% ethylene carbonate, 20% dextran sulfate, 600mM NaCl, 10mM sodium citrate (Matthiesen and Hansen, 2012). Slides were covered with parafilm, placed in humidified chambers, and allowed to incubate for 1 hour at 42^°^C. Slides were subsequently washed in 2X SSC at 37^°^C then allowed to air dry. Alexa Fluor 488-coupled tyramide signal amplification (Thermo Fisher Scientific T20912) was performed to increase probe signal above background autofluorescence from tick midgut digestive products (Biegala et al., 2002; Sojka et al., 2013). Slides were mounted in Diamond Prolong mounting medium plus DAPI and coverslips sealed with nail polish.

### Imaging

Confocal microscopy was performed at the University of Washington Keck Microscopy Center, on a Leica SP8X confocal microscope. 2-micron stacks were imaged at 63x magnification, using 6X averaging. Images were extracted from raw.lif files, maximum projected, channels merged, cropped, and pseudocolored using Fiji (Schindelin et al., 2012).

### Identification of interbacterial effector–immunity gene homologues in genomes

Multi-alignments for 220 putative and validated effector and immunity proteins were acquired from the supplemental materials of Zhang *et al*. (Zhang et al., 2012). Amino acid seed sequences of interbacterial effector and immunity proteins were queried via tBLASTn against a custom database of 63 *Spirochetes* genomes acquired from RefSeq, or a database of 22 genomes for tick-associated endosymbionts and pathogens from the phylum *Proteobacteria*. All hits with e-values less than or equal to 10^−3^ across the full length of the seed sequence were considered to be homologues of interbacterial effector or immunity proteins. The distribution of effector and immunity genes per genome within each genus was calculated by normalizing the total number of BLAST hit by the number of genomes searched.

## Author contributions

BDR, SC, and JDM conceived the study. BDR, SC, and JDM designed the study. BDR, SC, BH, and MCR conducted experimental work. XL, TJ, DN, and JB collected ticks. BDR, SC, and JDM wrote the paper. All authors read and approved the paper.

## Acknowledgements

We thank the Fred Hutch Experimental Histopathology Core facility and Dan Long at Rocky Mountain Labs for tick sectioning and slide preparation. We thank Barbara Simon, Jim Ruppa, David Simon, and June Reznikoff for their indefatigable efforts in collection of ticks from the Klickitat River “Ant Ranch”, and Susan Little for collection of Amblyomma ticks. We thank the UW Keck Imaging Center for providing equipment and assistance in confocal microscopy and acknowledge its support from the NIH (S10 OD016240). We thank Mr. DNA for sequencing support. We are grateful to colleagues for careful review of the manuscript and members of the Mougous lab for helpful discussions. This work was supported by National Institutes of Health grant R21AI114923 (JDM), and the Burroughs Wellcome Fund (JDM). BDR was supported by a Simons Foundation-sponsored Life Sciences Research Foundation postdoctoral fellowship.

## Data accessibility

Sequencing data was deposited at the NCBI SRA, BioProject accession # pending.

## Author information

The authors declare that no competing financial information exists. Correspondence should be directed to, and. 20

## Figure Legends

**Figure S1.**
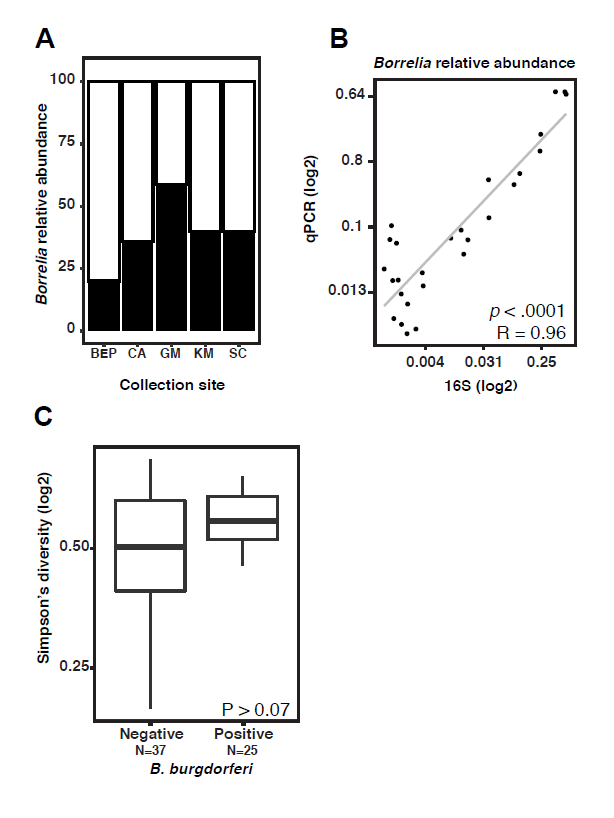
Characterization of *Borrelia burgdorferi* infection in wild *I. scapularis*. **(A)** *Borrelia* infection frequency as determined by qPCR for *B. burgdorferi flab* gene, plotted by collection site**. (B)** The relative abundance of *B. burgdorferi* as measured by qPCR with *flaB* primers calculated as a percent of total 16S rRNA gene counts exhibits a strong positive correlation between the relative abundance of *Borrelia* by 16S rRNA gene sequencing (Pearson *p* < 0.0001, *r* = 0.96). **(C)** Alpha diversity as quantified by the Simpson’s diversity metric, plotted for *B. burgdorferi* infected and uninfected ticks. No significant difference is observed (t-test, *p* > 0.07).

**Figure S2.**
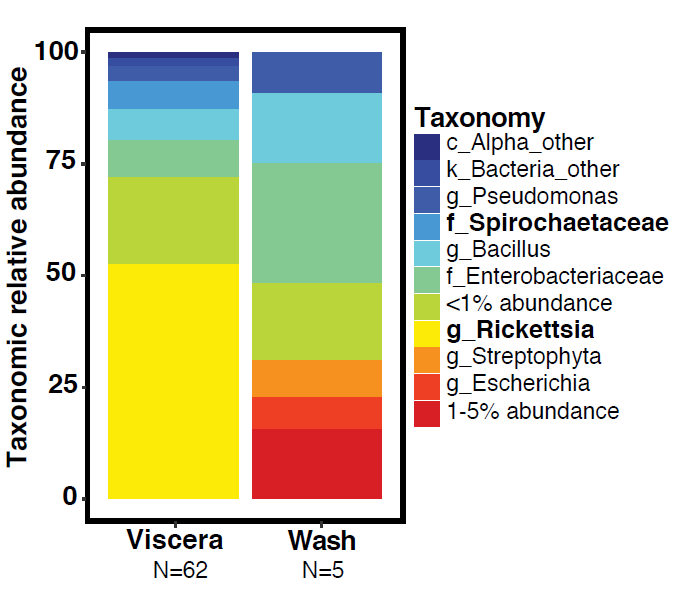
Taxonomic characterization of the *I. scapularis* microbiota. Relative taxonomic abundance for bacteria detected by 16S rRNA gene sequencing averaged across all adult *I. scapularis* internal viscera samples and external washes. Taxa previously known to be associated with *I. scapularis* are bolded in legend.

**Figure S3.**
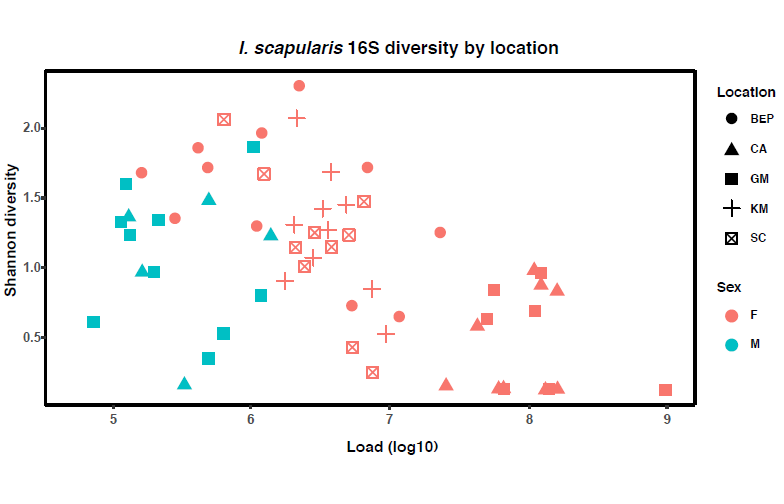
Bacterial load and diversity in adult female *I. scapularis* ticks across sample collection sites. Shannon diversity (y-axis) and bacterial load as measured by qPCR (x-axis) of adult female *I. scapularis* ticks. Samples cluster by sex, but not by geographic collection site. Big Eau Pleine (BEP), Carlos Avery (CA), Gordie Mikkelson (GM), Kettle Moraine (KM), Sandberg Conservancy (SC).

**Figure S4.**
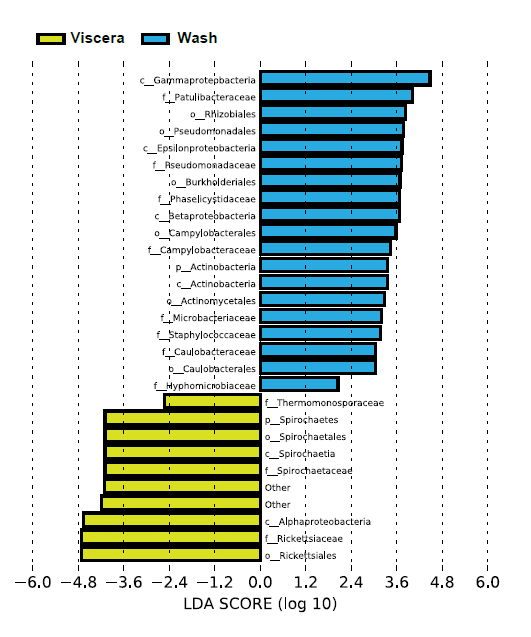
LEfSe plot of *I. scapularis* samples. Taxa enriched in either the viscera (yellow) or the wash samples (blue) as measured by the LDA score are indicated.

**Figure S5.**
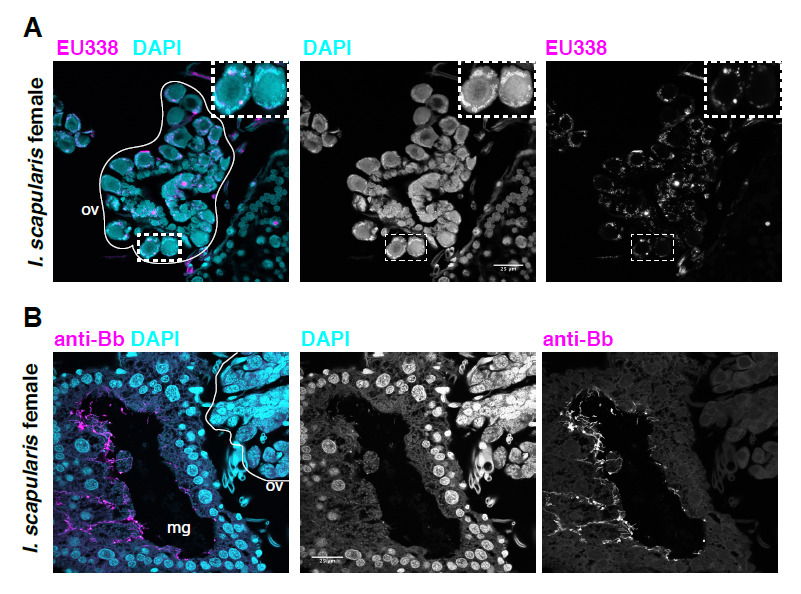
Detection of internal bacteria within *I. scapularis* by confocal microscopy. **A)** DAPI (blue) and EU338-TSA-ISH (magenta) staining of the central section of the ovary of a female adult *I. scapularis* tick. The ovary is outlined in white and region of inset zoom is indicated by a dashed white rectangle. **(B)** Anti-*B. burgdorferi* (magenta) and DAPI (blue) staining of a midgut cross-section and ovary of an infected *I. scapularis* adult female tick. The midgut lumen is indicated by “mg” and ovary is outlined in white. Scale bars are 25 microns.

**Figure S6.**
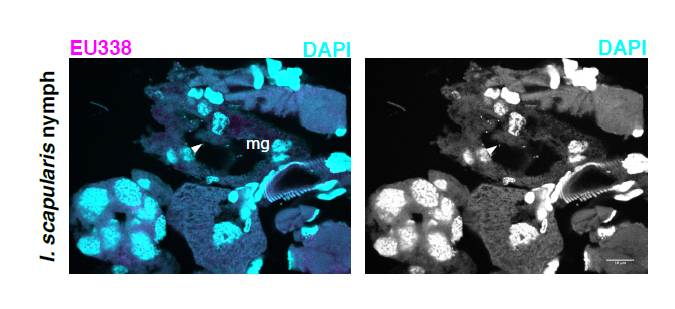
Confocal microscopy of *I. scapularis* nymphs. DAPI (blue) and EU338-TSA-ISH (magenta) staining of an *I. scapularis* nymph. Midgut indicated by arrowhead. Scale bar is 10 microns.

**Figure S7.**
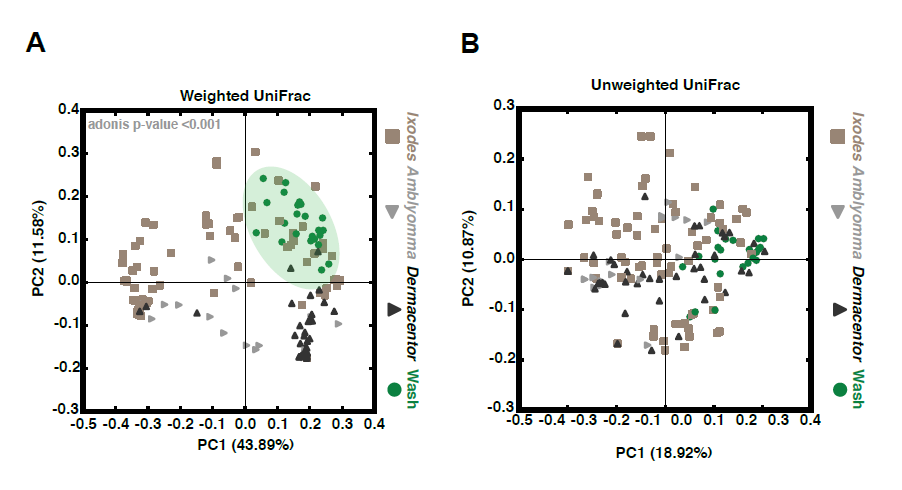
Beta diversity analysis of adult hard tick samples. **A)** Beta diversity analysis using weighted UniFrac (which accounts for taxon relative abundance within samples) reveals clustering by external wash samples and by tick species. **B)** Beta diversity analysis using the unweighted UniFrac metric (in which the relative abundance of taxa within samples is not taken into account) shows a lack of species-specific clustering.

**Figure S8.**
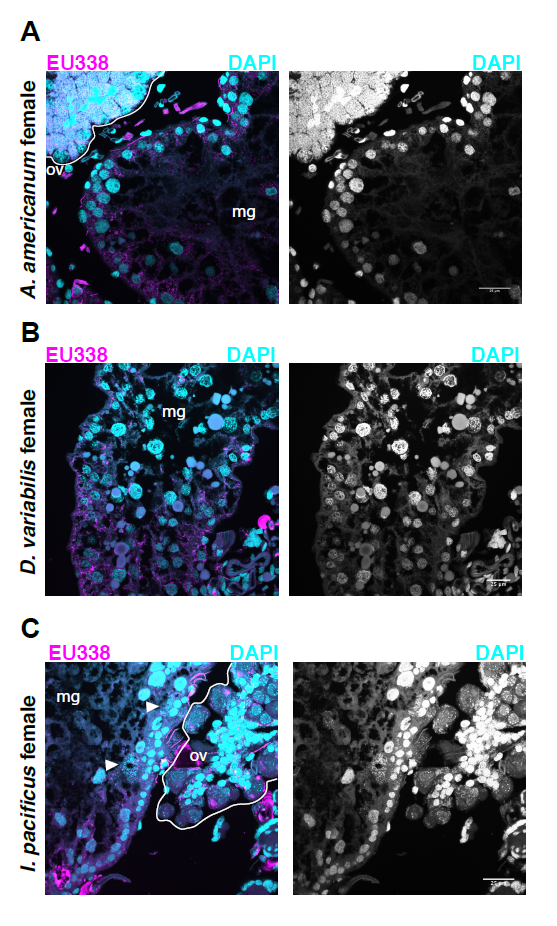
Confocal microscopy of non*-I. scapularis* hard ticks. **(A)** *Amblyomma maculatum* adult female, **(B)** *Dermacentor andersonii* adult female, **(C)** and *Ixodes pacificus* adult female. All ticks stained with DAPI (blue) and EU338-TSA-ISH (magenta), with the midgut lumen indicated with “mg” and ovary with “ov”. Arrowheads indicate bacterial micro-colonies within midgut. Ovaries are outlined in white in A and C. Scale bars are 25 microns.

## References

Abraham, N.M., Liu, L., Jutras, B.L., Yadav, A.K., Narasimhan, S., Gopalakrishnan, V., Ansari, J.M., Jefferson, K.K., Cava, F., Jacobs-Wagner, C., et al. (2017). Pathogen-mediated manipulation of arthropod microbiota to promote infection. Proc Natl Acad Sci U S A 114, E781–E790.

Amann, R.I., Ludwig, W., and Schleifer, K.H. (1995). Phylogenetic identification and in situ detection of individual microbial cells without cultivation. Microbiol Rev 59, 143–169.

Andreotti, R., Perez de Leon, A.A., Dowd, S.E., Guerrero, F.D., Bendele, K.G., and Scoles, G.A. (2011). Assessment of bacterial diversity in the cattle tick Rhipicephalus (Boophilus) microplus through tag-encoded pyrosequencing. BMC Microbiol 11, 6.

Anzaldi, L.L., and Skaar, E.P. (2010). Overcoming the heme paradox: heme toxicity and tolerance in bacterial pathogens. Infect Immun 78, 4977–4989.

Becker, N.S., Margos, G., Blum, H., Krebs, S., Graf, A., Lane, R.S., Castillo-Ramirez, S., Sing, A., and Fingerle, V. (2016). Recurrent evolution of host and vector association in bacteria of the Borrelia burgdorferi sensu lato species complex. BMC Genomics 17, 734.

Berende, A., ter Hofstede, H.J., Vos, F.J., van Middendorp, H., Vogelaar, M.L., Tromp, M., van den Hoogen, F.H., Donders, A.R., Evers, A.W., and Kullberg, B.J. (2016). Randomized Trial of Longer-Term Therapy for Symptoms Attributed to Lyme Disease. N Engl J Med 374, 1209–1220.

Biegala, I.C., Kennaway, G., Alverca, E., Lennon, J.-F., Vaulot, D., and Simon, N. (2002). Identification Of Bacteria Associated With Dinoflagellates (Dinophyceae) Alexandrium Spp. Using Tyramide Signal Amplification– Fluorescent In Situ Hybridization And Confocal Microscopy1. J Phycol 38, 404–411.

Budachetri, K., Browning, R.E., Adamson, S.W., Dowd, E., Chao, C.-C., Ching, W.-M., Karim, S., and Dowd, S.E. (2014). An Insight Into the Microbiome of the Amblyomma maculatum (Acari: Ixodidae) An Insight Into the Microbiome of the Amblyomma maculatum (Acari: Ixodidae). Journal of Medical Entomology 51, 119–129.

Budachetri, K., Williams, J., Mukherjee, N., Sellers, M., Moore, F., and Karim, S. (2016). The microbiome of neotropical ticks parasitizing on passerine migratory birds. Ticks Tick Borne Dis 8, 170–173.

Buffie, C.G., and Pamer, E.G. (2013). Microbiota-mediated colonization resistance against intestinal pathogens. Nat Rev Immunol 13, 790–801.

Cao, Z., Casabona, M.G., Kneuper, H., Chalmers, J.D., and Palmer, T. (2016). The type VII secretion system of Staphylococcus aureus secretes a nuclease toxin that targets competitor bacteria. Nat Microbiol 2, 16183.

Caporaso, J.G., Kuczynski, J., Stombaugh, J., Bittinger, K., Bushman, F.D., Costello, E.K., Fierer, N., Peña, A.G., Goodrich, J.K., Gordon, J.I., et al. (2010). QIIME allows analysis of high-throughput community sequencing data. Nat Methods 7, 335–336.

Chou, S., Daugherty, M.D., Peterson, S.B., Biboy, J., Yang, Y., Jutras, B.L., Fritz-Laylin, L.K., Ferrin, M.a., Harding, B.N., Jacobs-Wagner, C., et al. (2014). Transferred interbacterial antagonism genes augment eukaryotic innate immune function. Nature.

Cirimotich, C.M., Dong, Y., Clayton, A.M., Sandiford, S.L., Souza-Neto, J.A., Mulenga, M., and Dimopoulos, G. (2011a). Natural microbe-mediated refractoriness to Plasmodium infection in Anopheles gambiae. Science 332, 855–858.

Cirimotich, C.M., Ramirez, J.L., and Dimopoulos, G. (2011b). Native microbiota shape insect vector competence for human pathogens. Cell Host Microbe 10, 307–310.

Clay, K., Klyachko, O., Grindle, N., Civitello, D., Oleske, D., and Fuqua, C. (2008). Microbial communities and interactions in the lone star tick, Amblyomma americanum. Molecular Ecology 17, 4371–4381.

Clayton, K.A., Gall, C.A., Mason, K.L., Scoles, G.A., and Brayton, K.A. (2015). The characterization and manipulation of the bacterial microbiome of the Rocky Mountain wood tick, Dermacentor andersoni. Parasites & vectors 8, 632–632.

Durand, J., Herrmann, C., Genne, D., Sarr, A., Gern, L., and Voordouw, M.J. (2017). Multistrain Infections with Lyme Borreliosis Pathogens in the Tick Vector. Appl Environ Microbiol 83.

Durand, J., Jacquet, M., Paillard, L., Rais, O., Gern, L., and Voordouw, J. (2015). Cross-Immunity and Community Structure of a Multiple-Strain. 81, 7740–7752.

Dworkin, M.S., Schwan, T.G., and Anderson, D.E. Jr. (2002). Tick-borne relapsing fever in North America. Med Clin North Am 86, 417–433, viii-ix.

Edgar, R.C. (2010). Search and clustering orders of magnitude faster than BLAST. Bioinformatics 26, 2460–2461.

Edgar, R.C. (2013). UPARSE: highly accurate OTU sequences from microbial amplicon reads. Nat Methods 10, 996–998.

Finney, C.A., Kamhawi, S., and Wasmuth, J.D. (2015). Does the arthropod microbiota impact the establishment of vector-borne diseases in mammalian hosts? PLoS Pathog 11, e1004646.

Gall, C.A., Reif, K.E., Scoles, G.A., Mason, K.L., Mousel, M., Noh, S.M., and Brayton, K.A. (2016). The bacterial microbiome of Dermacentor andersoni ticks influences pathogen susceptibility. The ISME Journal, 1–10.

Gillespie, J.J., Joardar, V., Williams, K.P., Driscoll, T., Hostetler, J.B., Nordberg, E., Shukla, M., Walenz, B., Hill, C.a., Nene, V.M., et al. (2012). A rickettsia genome overrun by mobile genetic elements provides insight into the acquisition of genes characteristic of an obligate intracellular lifestyle. Journal of Bacteriology 194, 376–394.

Glassing, A., Dowd, S.E., Galandiuk, S., Davis, B., and Chiodini, R.J. (2016). Inherent bacterial DNA contamination of extraction and sequencing reagents may affect interpretation of microbiota in low bacterial biomass samples. Gut Pathog 8, 24.

Hamer, S.A., Hickling, G.J., Walker, E.D., and Tsao, J.I. (2014). Increased diversity of zoonotic pathogens and Borrelia burgdorferi strains in established versus incipient Ixodes scapularis populations across the Midwestern United States. Infection, Genetics and Evolution 27, 531–542.

Hammer, T.J., Janzen, D.H., Hallwachs, W., Jaffe, S.P., and Fierer, N. (2017). Caterpillars lack a resident gut microbiome. Proc Natl Acad Sci U S A 114, 9641–9646.

Harvell, C.D., Mitchell, C.E., Ward, J.R., Altizer, S., Dobson, A.P., Ostfeld, R.S., and Samuel, M.D. (2002). Climate warming and disease risks for terrestrial and marine biota. Science 296, 2158–2162.

Hawlena, H., Rynkiewicz, E., Toh, E., Alfred, A., Durden, L.a., Hastriter, M.W., Nelson, D.E., Rong, R., Munro, D., Dong, Q., et al. (2012). The arthropod, but not the vertebrate host or its environment, dictates bacterial community composition of fleas and ticks. The ISME Journal 7, 221–223.

Hayes, C.S., Aoki, S.K., and Low, D.A. (2010). Bacterial contact-dependent delivery systems. Annu Rev Genet 44, 71–90.

Herrmann, C., Gern, L., and Voordouw, M.J. (2013). Species co-occurrence patterns among Lyme borreliosis pathogens in the tick vector Ixodes ricinus. Applied and environmental microbiology 79, 7273–7280.

Hibbing, M.E., Fuqua, C., Parsek, M.R., and Peterson, S.B. (2010). Bacterial competition: surviving and thriving in the microbial jungle. Nature Reviews Microbiology 8, 15–25.

Ismail, N., Bloch, K.C., and McBride, J.W. (2010). Human ehrlichiosis and anaplasmosis. Clin Lab Med 30, 261–292.

Jervis-Bardy, J., Leong, L.E., Marri, S., Smith, R.J., Choo, J.M., Smith-Vaughan, H.C., Nosworthy, E., Morris, P.S., O’Leary, S., Rogers, G.B., et al. (2015). Deriving accurate microbiota profiles from human samples with low bacterial content through post-sequencing processing of Illumina MiSeq data. Microbiome 3, 19.

Jewett, M.W., Lawrence, K., Bestor, A.C., Tilly, K., Grimm, D., Shaw, P., VanRaden, M., Gherardini, F., and Rosa, P.A. (2007). The critical role of the linear plasmid lp36 in the infectious cycle of Borrelia burgdorferi. Mol Microbiol 64, 1358–1374.

Jing, X., Wong, A.C., Chaston, J.M., Colvin, J., McKenzie, C.L., and Douglas, A.E. (2014). The bacterial communities in plant phloem-sap-feeding insects. Mol Ecol 23, 1433–1444.

Jones, B.A., Grace, D., Kock, R., Alonso, S., Rushton, J., Said, M.Y., McKeever, D., Mutua, F., Young, J., McDermott, J., et al. (2013). Zoonosis emergence linked to agricultural intensification and environmental change. Proc Natl Acad Sci U S A 110, 8399–8404.

Keesing, F., Belden, L.K., Daszak, P., Dobson, A., Harvell, C.D., Holt, R.D., Hudson, P., Jolles, A., Jones, K.E., Mitchell, C.E., et al. (2010). Impacts of biodiversity on the emergence and transmission of infectious diseases. Nature 468, 647–652.

Khoo, J.J., Chen, F., Kho, K.L., Ahmad Shanizza, A.I., Lim, F.S., Tan, K.K., Chang, L.Y., and AbuBakar, S. (2016). Bacterial community in Haemaphysalis ticks of domesticated animals from the Orang Asli communities in Malaysia. Ticks Tick Borne Dis 7, 929–937.

Kim, D., Hofstaedter, C.E., Zhao, C., Mattei, L., Tanes, C., Clarke, E., Lauder, A., Sherrill-Mix, S., Chehoud, C., Kelsen, J., et al. (2017). Optimizing methods and dodging pitfalls in microbiome research. Microbiome 5, 52.

Klyachko, O., Stein, B.D., Grindle, N., Clay, K., and Fuqua, C. (2007). Localization and visualization of a Coxiella-type symbiont within the lone star tick, Amblyomma americanum. Applied and Environmental Microbiology 73, 6584–6594.

Kocan, K.M., de la Fuente, J., Blouin, E.F., Coetzee, J.F., and Ewing, S.A. (2010). The natural history of Anaplasma marginale. Vet Parasitol 167, 95–107.

Kommineni, S., Bretl, D.J., Lam, V., Chakraborty, R., Hayward, M., Simpson, P., Cao, Y., Bousounis, P., Kristich, C.J., and Salzman, N.H. (2015). Bacteriocin production augments niche competition by enterococci in the mammalian gastrointestinal tract. Nature 526, 719–722.

Kugeler, K.J., Farley, G.M., Forrester, J.D., and Mead, P.S. (2015). Geographic Distribution and Expansion of Human Lyme Disease, United States. Emerg Infect Dis 21, 1455–1457.

Lauder, A.P., Roche, A.M., Sherrill-Mix, S., Bailey, A., Laughlin, A.L., Bittinger, K., Leite, R., Elovitz, M.A., Parry, S., Bushman, F.D., et al. (2016). Comparison of placenta samples with contamination controls does not provide evidence for a distinct placenta microbiota. Microbiome 4, 29–29.

Lescot, M., Audic, S., Robert, C., Nguyen, T.T., Blanc, G., Cutler, S.J., Wincker, P., Couloux, A., Claverie, J.M., Raoult, D., et al. (2008). The genome of Borrelia recurrentis, the agent of deadly louse-borne relapsing fever, is a degraded subset of tick-borne Borrelia duttonii. PLoS Genet 4, e1000185.

Matthiesen, S.H., and Hansen, C.M. (2012). Fast and non-toxic in situ hybridization without blocking of repetitive sequences. PLoS One 7, e40675.

Moreno, C.X., Moy, F., Daniels, T.J., Godfrey, H.P., and Cabello, F.C. (2006). Molecular analysis of microbial communities identified in different developmental stages of Ixodes scapularis ticks from Westchester and Dutchess Counties, New York. Environmental Microbiology 8, 761–772.

Nadkarni, M.A., Martin, F.E., Jacques, N.A., and Hunter, N. (2002). Determination of bacterial load by real-time PCR using a broad-range (universal) probe and primers set. Microbiology 148, 257–266.

Nakao, R., Abe, T., Nijhof, a.M., Yamamoto, S., Jongejan, F., Ikemura, T., and Sugimoto, C. (2013). A novel approach, based on BLSOMs (Batch Learning Self-Organizing Maps), to the microbiome analysis of ticks. ISME Journal 7, 1003–1015.

Narasimhan, S., and Fikrig, E. (2015). Tick microbiome: the force within. Trends in Parasitology 31, 315–323.

Narasimhan, S., Rajeevan, N., Liu, L., Zhao, Y.O., Heisig, J., Pan, J., Eppler-Epstein, R., Deponte, K., Fish, D., and Fikrig, E. (2014). Gut microbiota of the tick vector Ixodes scapularis modulate colonization of the Lyme disease spirochete. Cell host & microbe 15, 58–71.

Nigrovic, L.E., and Thompson, K.M. (2007). The Lyme vaccine: a cautionary tale. Epidemiol Infect 135, 1–8.

Noda, H., Munderloh, U.G., and Kurtti, T.J. (1997). Endosymbionts of ticks and their relationship to Wolbachia spp. and tick-borne pathogens of humans and animals. Appl Environ Microbiol 63, 3926–3932.

Palmer, W.J., and Jiggins, F.M. (2015). Comparative Genomics Reveals the Origins and Diversity of Arthropod Immune Systems. Mol Biol Evol 32, 2111–2129.

Paster, B.J., Dewhirst, F.E., Weisburg, W.G., Tordoff, L.A., Fraser, G.J., Hespell, R.B., Stanton, T.B., Zablen, L., Mandelco, L., and Woese, C.R. (1991). Phylogenetic analysis of the spirochetes. J Bacteriol 173, 6101–6109.

R Development Core Team (2013). R: A Language and Environment for Statistical Computing.

Radolf, J.D., Caimano, M.J., Stevenson, B., and Hu, L.T. (2012). Of ticks, mice and men: understanding the dual-host lifestyle of Lyme disease spirochaetes. Nature reviews Microbiology 10, 87–99.

Riley, M.A., and Wertz, J.E. (2002). Bacteriocins: evolution, ecology, and application. Annu Rev Microbiol 56, 117–137.

Russell, A.B., Peterson, S.B., and Mougous, J. (2014). Type VI secretion system effectors: poisons with a purpose. Nat Rev Microbiol 12, 137–148.

Rynkiewicz, E.C., Hemmerich, C., Rusch, D.B., Fuqua, C., and Clay, K. (2015). Concordance of bacterial communities of two tick species and blood of their shared rodent host. Molecular Ecology 24, 2566–2579.

Salter, S.J., Cox, M.J., Turek, E.M., Calus, S.T., Cookson, W.O., Moffatt, M.F., Turner, P., Parkhill, J., Loman, N.J., and Walker, A.W. (2014). Reagent and laboratory contamination can critically impact sequence-based microbiome analyses. BMC Biology 12, 87–87.

Schindelin, J., Arganda-Carreras, I., Frise, E., Kaynig, V., Longair, M., Pietzsch, T., Preibisch, S., Rueden, C., Saalfeld, S., Schmid, B., et al. (2012). Fiji: an open-source platform for biological-image analysis. Nat Methods 9, 676–682.

Segata, N., Izard, J., Waldron, L., Gevers, D., Miropolsky, L., Garrett, W.S., and Huttenhower, C. (2011). Metagenomic biomarker discovery and explanation. Genome Biol 12, R60.

Simhadri, R.K., Fast, E.M., Guo, R., Schultz, M.J., Vaisman, N., Ortiz, L., Bybee, J., Slatko, B.E., and Frydman, H.M. (2017). The Gut Commensal Microbiome of Drosophila melanogaster Is Modified by the Endosymbiont Wolbachia. mSphere 2.

Sojka, D., Franta, Z., Horn, M., Caffrey, C.R., Mareš, M., and Kopáček, P. (2013). New insights into the machinery of blood digestion by ticks. Trends Parasitol 29, 276–285.

Sonenshine, D.E., and Roe, R.M. (2014). Biology of ticks, 2nd edn (New York: Oxford University Press).

Souza, D.P., Oka, G.U., Alvarez-Martinez, C.E., Bisson-Filho, A.W., Dunger, G., Hobeika, L., Cavalcante, N.S., Alegria, M.C., Barbosa, L.R., Salinas, R.K., et al. (2015). Bacterial killing via a type IV secretion system. Nat Commun 6, 6453.

Steinhaus, E.A. (1946). Insect microbiology; an account of the microbes associated with insects and ticks, with special reference to the biologic relationships involved (Ithaca, N.Y.,: Comstock publishing company, inc.).

Strandh, M., and Råberg, L. (2015). Within-host competition between Borrelia afzelii ospC strains in wild hosts as revealed by massively parallel amplicon sequencing. Philosophical transactions of the Royal Society of London Series B, Biological sciences 370, 20140293–20140293-.

Swei, A., and Kwan, J.Y. (2016). Tick microbiome and pathogen acquisition altered by host blood meal. The ISME Journal, 1–4.

Trout Fryxell R.T., and DeBruyn, J.M. (2016). The Microbiome of Ehrlichia-Infected and Uninfected Lone Star Ticks (Amblyomma americanum). PLoS One 11, e0146651.

van Treuren, W., Ponnusamy, L., Brinkerhoff, R.J., Gonzalez, A., Parobek, C.M., Juliano, J.J., Andreadis, T.G., Falco, R.C., Ziegler, L.B., Hathaway, N., et al. (2015). Variation in the microbiota of Ixodes ticks with regard to geography, species, and sex. Applied and Environmental Microbiology 81, 6200–6209.

Vannini, C., Petroni, G., Verni, F., and Rosati, G. (2005). A bacterium belonging to the Rickettsiaceae family inhabits the cytoplasm of the marine ciliate Diophrys appendiculata (Ciliophora, Hypotrichia). Microbial Ecology 49, 434–442.

Verster, A.J., Ross, B.D., Radey, M.C., Bao, Y., Goodman, A.L., Mougous, J.D., and Borenstein, E. (2017). The Landscape of Type VI Secretion across Human Gut Microbiomes Reveals Its Role in Community Composition. Cell Host Microbe 22, 411–419 e414.

Voordouw, M.J. (2015). Co-feeding transmission in Lyme disease pathogens. Parasitology 142, 290–302.

Vora, N. (2008). Impact of anthropogenic environmental alterations on vector-borne diseases. Medscape J Med 10, 238.

Wang, S., Dos-Santos, A.L.A., Huang, W., Liu, K.C., Oshaghi, M.A., Wei, G., Agre, P., and Jacobs-Lorena, M. (2017). Driving mosquito refractoriness to Plasmodium falciparum with engineered symbiotic bacteria. Science 357, 1399.

Wexler, A.G., Bao, Y., Whitney, J.C., Bobay, L.-M., Xavier, J.B., Schofield, W.B., Barry, N.A., Russell, A.B., Tran, B.Q., Goo, Y.A., et al. (2016). Human symbionts inject and neutralize antibacterial toxins to persist in the gut. Proc Natl Acad Sci U S A 113, 3639–3644.

Whitney, J.C., Peterson, S.B., Kim, J., Pazos, M., Verster, A.J., Radey, M.C., Kulasekara, H.D., Ching, M.Q., Bullen, N.P., Bryant, D., et al. (2017). A broadly distributed toxin family mediates contact-dependent antagonism between gram-positive bacteria. eLife 6.

Williams-Newkirk, A.J., Rowe, L.A., Mixson-Hayden, T.R., and Dasch, G.A. (2014). Characterization of the bacterial communities of life stages of free living lone star ticks (Amblyomma americanum). PLoS ONE 9.

Wormser, G.P., Dattwyler, R.J., Shapiro, E.D., Halperin, J.J., Steere, A.C., Klempner, M.S., Krause, P.J., Bakken, J.S., Strle, F., Stanek, G., et al. (2006). The Clinical Assessment, Treatment, and Prevention of Lyme Disease, Human Granulocytic Anaplasmosis, and Babesiosis: Clinical Practice Guidelines by the Infectious Diseases Society of America. Clinical Infectious Diseases 43, 1089–1134.

Zhang, D., de Souza, R.F., Anantharaman, V., Iyer, L.M., and Aravind, L. (2012). Polymorphic toxin systems: Comprehensive characterization of trafficking modes, processing, mechanisms of action, immunity and ecology using comparative genomics. Biol Direct 7, 18–18.

Zhang, K., Bian, J., Deng, Y., Smith, A., Nunez, R.E., Li, M.B., Pal, U., Yu, A.M., Qiu, W., Ealick, S.E., et al. (2016). Lyme disease spirochaete Borrelia burgdorferi does not require thiamin. Nat Microbiol 2, 16213.

Zolnik, C.P., Prill, R.J., Falco, R.C., Daniels, T.J., and Kolokotronis, S.-O. (2016). Microbiome changes through ontogeny of a tick pathogen vector. Molecular Ecology.

